# The determinants of the rarity of nucleic and peptide short sequences in nature

**DOI:** 10.1101/2023.09.24.559219

**Authors:** Nikol Chantzi, Ioannis Mouratidis, Manvita Mareboina, Maxwell A. Konnaris, Austin Montgomery, Ilias Georgakopoulos-Soares

## Abstract

The prevalence of nucleic and peptide short sequences across organismal genomes and proteomes has not been thoroughly investigated. Here we examined 45,785 reference genomes and 21,871 reference proteomes, spanning archaea, bacteria, viruses and eukaryotes to calculate the rarity of short sequences in them. To capture this, we developed a metric of the rarity of each sequence in nature, the Anti-Kardashian index. We find that the frequency of certain dipeptides in rare oligopeptide sequences is hundreds of times lower than expected, which is not the case for any dinucleotides. We also generate predictive regression models that infer the rarity of nucleic and proteomic sequences in nature. For six-mer peptide kmers the R^2^ performance of the regression models based on amino acid and dipeptide content is 0.816, whereas models based on physicochemical features achieve an R^2^ of 0.788. For twelve-mer nucleic kmers the R^2^ performance of our models based on mono and dinucleotides is 0.481. Our results indicate that the mono and dinucleotide composition of nucleic sequences and the amino acids, dipeptides and physicochemical properties of peptide sequences can explain a significant proportion of the variance in their frequencies between organisms in nature.

## Introduction

Genomic and proteomic information is often measured using kmers, which are contiguous sequences of length *k* composed of nucleotides in genomics or amino acids in proteomics. Kmers are distributed inhomogeneously in DNA and protein molecules. The frequency spectrum of kmers can be analyzed, which displays the distribution of kmer frequencies in a single genome or proteome (Yang et al. 2020; Chor et al. 2009; Mittal, Changani, and Taparia 2020). At the genomic level, the GC and CpG content influence the frequency of kmer sequences (Chae et al. 2013). Other features, such as *cis*-regulatory elements, constraint genomic regions and pathogenic variant sites also display specific kmer preferences and shape the kmer spectrum (di Iulio et al. 2018). At the proteomic level, the energy expenditure associated with each amino acid influences the kmer peptide frequency spectrum (Swire 2007). In addition, specific functional sites and common protein motifs account for overrepresented kmers (Poznański et al. 2018).

There are also kmer sequences that are rare or absent from one or more genomes or proteomes. Kmer sequences that are absent from a genome or proteome are referred to as nullomers and nullpeptides, respectively (Hampikian and Andersen 2007; Tuller, Chor, and Nelson 2007; Fofanov et al. 2004). The absence of kmer sequences has been previously attributed to selection constraints and hypermutation in nucleic sequences and on structural and chemical constraints for peptide sequences (Vergni and Santoni 2016; Georgakopoulos-Soares et al. 2021; Koulouras and Frith 2021). Kmers absent from every genome or proteome are referred to as primes (Hampikian and Andersen 2007; Georgakopoulos-Soares et al. 2021). Additionally, quasi-primes were recently proposed as the shortest kmer sequences, which are unique to a species’s genome or proteome and absent from every other organism (Mouratidis et al. 2023). As an extension, certain kmer sequences can be preferentially found within specific phylogenetic groups and a subset of kmers are taxonomy-specific (Mouratidis et al. 2023). However, none of the existing approaches provides a measure for the rarity of every nucleic or peptide sequence and therefore such an index of kmer rarity across organisms is lacking.

Here we introduce the Anti-Kardashian index, a measure of anti-popularity of genomic or proteomic sequences in nature (**Figure 1**). To calculate the rarity of kmer sequences we examine 45,785 reference genomes and 21,871 reference proteomes, spanning archaea, bacteria, viruses and eukaryotes, from which we extract the set of kmers in each of them, to estimate the rarity of individual kmers, within and across taxonomies. We find that there are significant compositional biases in rare sequences, both in peptide and nucleic kmers. We generate predictive models that estimate the rarity of each kmer sequence, trained on the primary mono- and di-nucleotide content of nucleic kmers or using the amino acid and di-peptide content of each peptide kmer. Additionally, we generate models trained on physicochemical information of each peptide kmer, that achieve comparable results. These findings provide evidence that the rarity of biological information in nature can be inferred.

**Figure 1:**
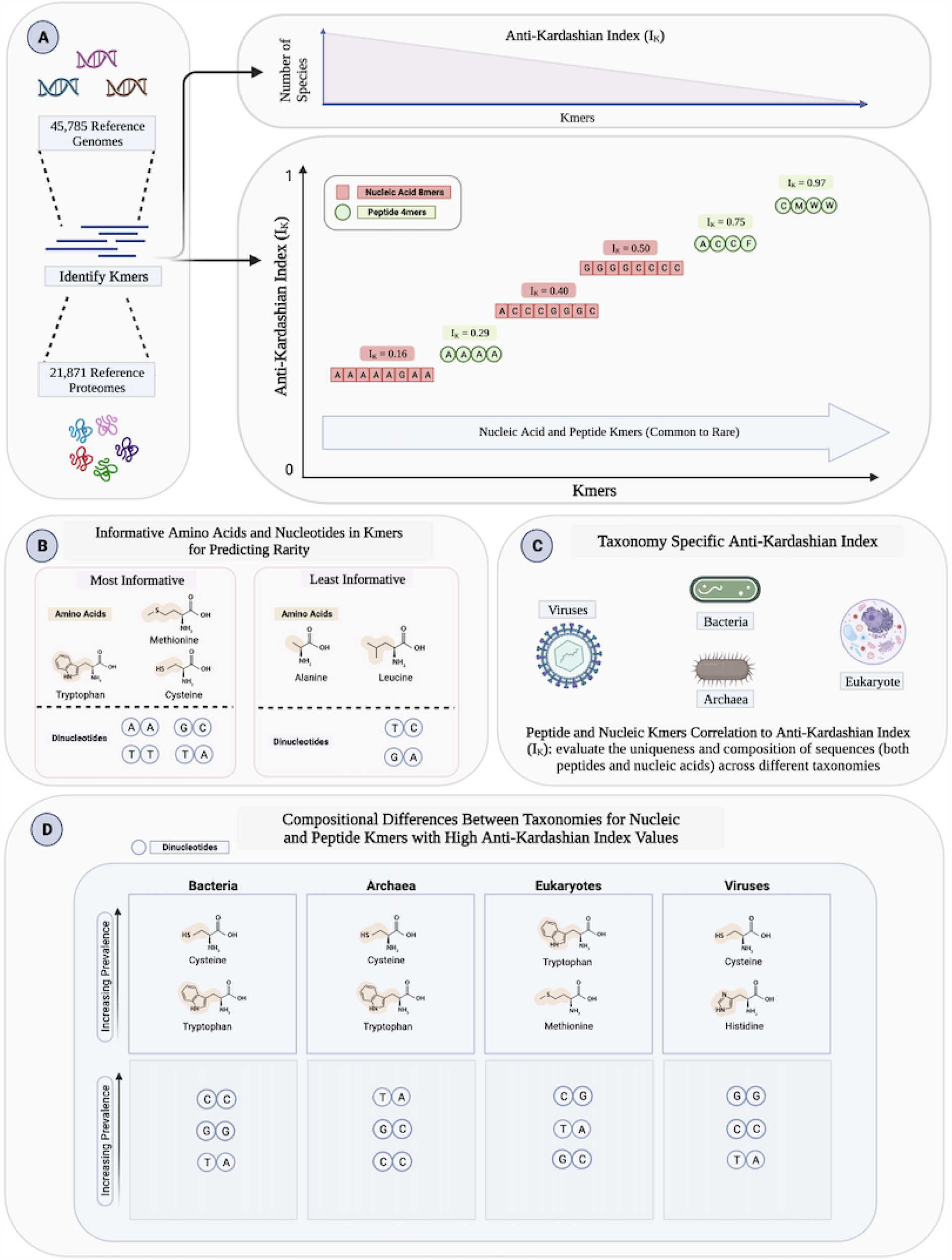
Schematic illustration of the Anti-Kardashian index, an estimate of the rarity of each kmer peptide or nucleic sequence in nature. **A**. Schematic displaying the identification of Anti-Kardashian index to depict the rarity of various peptide and nucleic acid kmers. All kmers were identified from 45, 785 reference genomes and 21, 871 reference proteomes. Nucleic acid and peptide kmer Anti-Kardashian Index values were identified on a scale from 0 to 1, with values closer to 1 indicating increased sequence absence across organisms. **B**. Schematic shows the various amino acids and nucleotides in kmers that predict rarity, specifying the most and least informative amino acids and dinucleotides. **C**. Taxonomy-specific peptide and nucleic kmers were determined to evaluate the uniqueness of sequences across various organismal groups. **D**. Compositional differences across taxonomies were depicted to show that the decreasing prevalence of kmers produces an increasing prevalence of these associated dinucleotides and peptides.

## Results

### Derivation of the Anti-Kardashian index

Given an alphabet A and a string consisting of kmers drawn from that alphabet - under the assumption that each letter is equiprobable and appears independently, as the string gets arbitrarily large, the probability of observing one specific kmer sequence diminishes substantially according to the following exponential law:

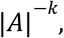

where k is the length of the string, and |*A*| the total number of letters of the alphabet. Mathematically, this means that as the kmer strings populate different species, if the length gets sufficiently large, the intersection would converge towards the empty set. Nonetheless, kmer sequences that consist of amino acids or nucleic acids are not random nor independent from one another, and, as such, they exhibit different distributional characteristics.

A kmer sequence can be present in a set of genomes or proteomes, and its presence can reflect phylogenetic relationships and evolutionary history, biological mechanisms and functions or can be stochastic. Here, we define the Anti-Kardashian index as a measure of the rarity of each kmer sequence across the examined organisms.

Mathematically the Anti-Kardashian index is defined below:

Let us define the set of all kmers of length *k* present in a genome or proteome, represented by *S*_*k*_. Furthermore, we define *T =* {*S*_1_, *S*_2_, … *S*_*n*_} as the set containing the sets of kmers present in each genome or proteome in our reference database. The Anti-Kardashian index *I*_*K*_ of a kmer K is defined as:

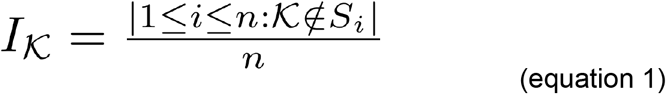

The Anti-Kardashian index *I*_*K*_ ranges between zero and one, with zero indicating a sequence found in every organism and one indicating a sequence absent across every species. The concept applies to different categories of biological sequences, including kmer peptides and nucleic acids.

### Frequency patterns of kmers across species

Using the set of 21,871 reference proteomes we estimated the number of species each kmer peptide was found in, for peptide kmer lengths between two and six amino acids. The kmer length limit of six amino acids was used because at longer kmer lengths most peptides are absent from most proteomes (**Supplementary Figure 1**). We also investigated 45,785 reference genomes and estimated the frequency of each nucleic kmer across the examined species for kmer lengths between six and twelve base-pairs (bps). The kmer length limit of twelve bps was used because at lower kmer lengths the majority of genomes contain the majority of the possible kmers, not allowing sufficient differentiation of the species (**Supplementary Figure 2**). We observe that the number of kmers detected per species is highest across eukaryotes and lowest across viruses, across the kmer lengths studied, when examining proteomes (**Supplementary Figure 1**) and genomes (**Supplementary Figure 2**). For instance, in six-mer peptides there are on average 5,090,888, 1,070,044, 581,726 and 12,026 kmers in eukaryotic, bacterial, archaeal and viral proteomes, respectively (**Supplementary Figure 1**). Furthermore, in twelve-mer nucleic kmers there are 14,248,979, 4,444,638, 3,225,470 and 83,603 different kmers in eukaryotic, bacterial, archaea and viral genomes (**Supplementary Figure 2**).

Next, for each kmer peptide we estimated its Anti-Kardashian index (*I*_*K*_), as a measure of the rarity of individual kmer peptide sequences in nature. We observe that the Anti-Kardashian index of the kmer peptides increased as a function of kmer length and shifted from zero to one (Pearson correlation = 0.991, p-value <0.0001; **Figure 2a-b**; **Supplementary Figure 3a**). We also observe that after four amino acids kmer length, the majority of kmers are absent from most proteomes, while the first peptide prime sequences, which are absent from every proteome, appear at six amino acids kmer length, having an Anti-Kardashian index of one.

**Figure 2:**
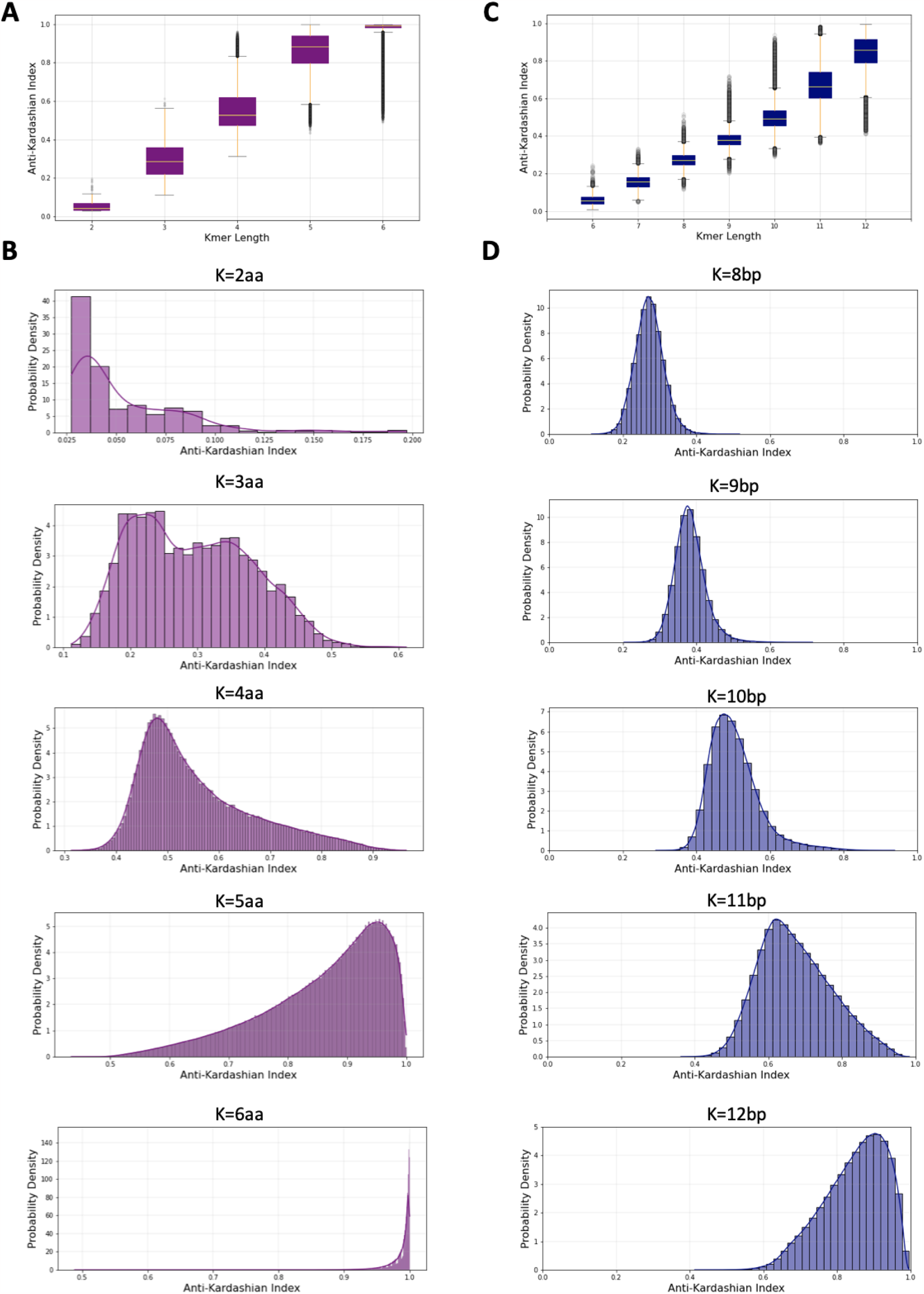
Oligopeptide and oligonucleotide Anti-Kardashian kmer profiles. **A**. The Anti-Kardashian index as a function of peptide kmer length. **B**. Probability density for the Anti-kardashian kmer profile for oligopeptide lengths of two to seven amino acids. **C**. The Anti-Kardashian index as a function of nucleic kmer length. **D**. Probability density for the Anti-kardashian kmer profile for oligonucleotide lengths of six to twelve base pairs. An Anti-Kardashian index of zero indicates that a sequence is found in every species and an Anti-Kardashian index of one indicates that a sequence is absent across all species.

From the frequency of each kmer across the reference genomes we derived the Anti-Kardashian index score of each nucleic kmer. Similarly to the proteome analysis, we find that the number of species each kmer appears in, decreases as a function of kmer length, and the Anti-Kardashian index distribution converges towards one (Pearson correlation = 0.992, p-value <1.3e-05; **Figure 2c-d**; **Supplementary Figure 3b**). These results also indicate that, with the exception of eukaryotes, after ten bps length, most potential kmers are absent from a genome and that longer kmers have a higher Anti-Kardashian index across sequence types. Additionally, a small subset of nucleic and peptide kmers remains highly frequent, even at longer kmer lengths, resulting in a negative skew with a long tail distribution (**Figure 2 b,d**), which likely reflects repetitive or biologically functional kmer sequences.

### Rare peptide and nucleic sequences have a distinct sequence composition

Furthermore, we examined the Anti-Kardashian index of peptide and nucleic kmers across each species, spanning the three domains of life and in viruses. For peptide sequences, we find that clustering species based on the Anti-Kardashian index score of each of the constituent kmers separates the taxonomies (**Figure 3a**). We also observe that the taxonomic clustering performance is influenced by the kmer length, with longer peptide kmers showing improved clustering of taxonomic sub-groups (**Figure 3a**; **Supplementary Figure 4**). The improved clustering between taxonomic groups at longer kmer lengths reflects the higher likelihood of long kmer sequences being found in fewer, and more closely related organismal proteomes.

**Figure 3:**
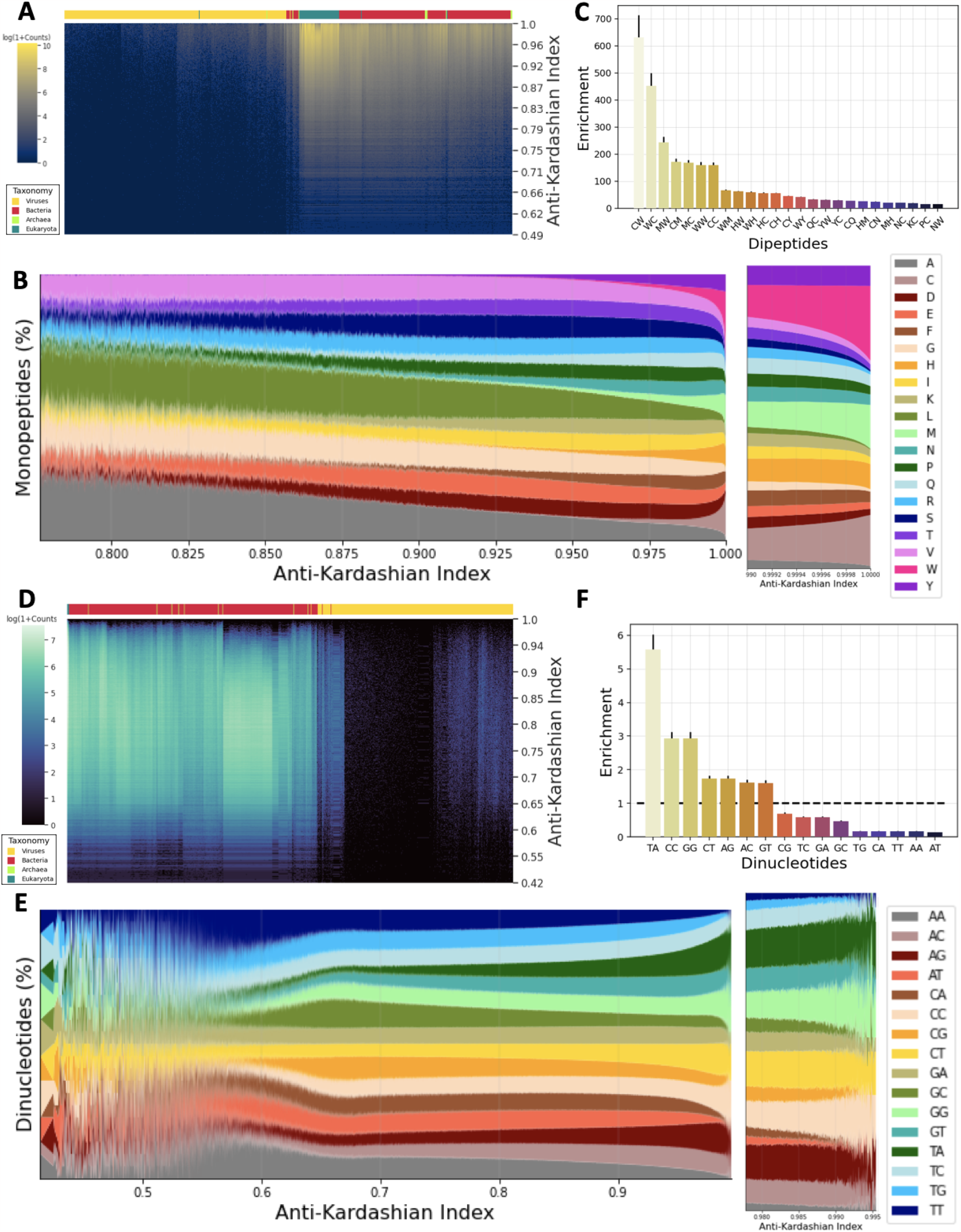
The Anti-Kardashian index scores across kmers in proteomes and genomes and across the taxonomic subgroups. **A**. Kmers detected in each species as a function of the Anti-Kardashian index score for six-mer peptide kmers. Color bar indicates the taxonomic group, namely viruses, bacteria, archaea and eukaryotes. **B**. Association between the Anti-Kardashian Index and the amino acid and dipeptide content of kmers in proteomes and genomes. **C**. Enrichment of dipeptides for frequencies in kmer sequences with Anti-Kardashian index>0.95 over sequences with Anti-Kardashian index<0.9, for kmer length of five amino acids. **D**. Kmers detected in each species as a function of the Anti-Kardashian index score for twelve bps kmer length. Color bar indicates the taxonomic group, namely viruses, bacteria, archaea and eukaryotes. **E**. The frequency of each amino acid is estimated for different Anti-Kardashian index scores. Color bar indicates each amino acid. **E**. The frequency of each dinucleotide is estimated for different Anti-Kardashian index scores. Color bar indicates each dinucleotide. **F**. Enrichment of dinucleotides for frequencies in kmer sequences with Anti-Kardashian index>0.99 over sequences with Anti-Kardashian index<0.5, for twelve bps kmer length.

We were interested to investigate if the rarity of kmer sequences can be explained by their sequence composition. We therefore examined the amino acid and dinucleotide content of peptide and nucleic kmers as a function of the Anti-Kardashian index. We find that for peptides, tryptophan (W), methionine (M) and cysteine (C) increase in proportion for kmers of higher Anti-Kardashian index, whereas alanine and leucine disappear at the kmers with the highest Anti-Kardashian index across the kmer lengths studied (**Figure 3b**; **Supplementary Figure 5**). In particular, above the 99^th^ percentile of the Anti-Kardashian index, which reflects the rarest peptide sequences, tryptophan, methionine and cysteine increase precipitously and capture a significant proportion of the amino acid space. At dipeptides the bias is even greater, with CW, WC and MW having a 631-fold, 452-fold and 245-fold enrichment respectively in the frequency for kmers with an Anti-Kardashian index greater than 0.95 over sequences with an Anti-Kardashian index less than 0.9 (**Figure 3c**). These results indicate that rare or absent sequences are more likely to have specific amino acid or dipeptide content.

When clustering based on the Anti-Kardashian index of each kmer across the organismal reference genomes we also observe taxonomic clustering (**Figure 3d**; **Supplementary Figure 6**). We also investigate the frequencies of dinucleotides in nucleic kmers as a function of their Anti-Kardashian index and observe significant differences across the continuum. Increased frequencies of TA and GG/CC kmers are associated with higher Anti-Kardashian index (**Figure 3e**; **Supplementary Figure 7**). In contrast, AA/TT are enriched at kmers with low Anti-Kardashian index, which is particularly evident above the 99^th^ percentile across kmer lengths (**Figure 3e**; **Supplementary Figure 7**). For instance, for TA and CC/GG we observe 5.58-fold and 2.93 enrichments for twelve bp kmers with an Anti-Kardashian index greater than 0.99 over sequences with an Anti-Kardashian index less than 0.5 (**Figure 3f**). These results indicate differences in the nucleotide composition of rare nucleic kmers, which however are less pronounced than the differences observed for rare peptide kmers.

### Comparison of kmer frequency patterns between species of different taxonomies

We next estimated the Anti-Kardashian index of peptide and nucleic kmers within individual taxonomies, namely archaea, bacteria, eukaryotes and viruses, and examined the correlation of the taxonomic Anti-Kardashian index between them. Overall, we observe significant positive correlations between all the taxonomies both for peptide and nucleic kmers (**Figure 4a-b**). Specifically, for both peptide and nucleic kmers, the comparison of the Anti-Kardashian index between taxonomic groups revealed positive correlations between all taxonomies, with the viruses displaying the weakest correlations with the other taxonomic groups (**Figure 4a-b**). The strongest correlations were observed between archaea and bacteria (**Figure 4a-b**) and horizontal gene transfer could account for the higher degree of similarity. We also observe significant deviations for specific kmers for their taxonomic Anti-Kardashian index, when performing pairwise comparisons between the three domains of life and viruses (**Figure 4c-d**), which could reflect compositional differences between the taxonomies but also functional differences of biological importance.

**Figure 4:**
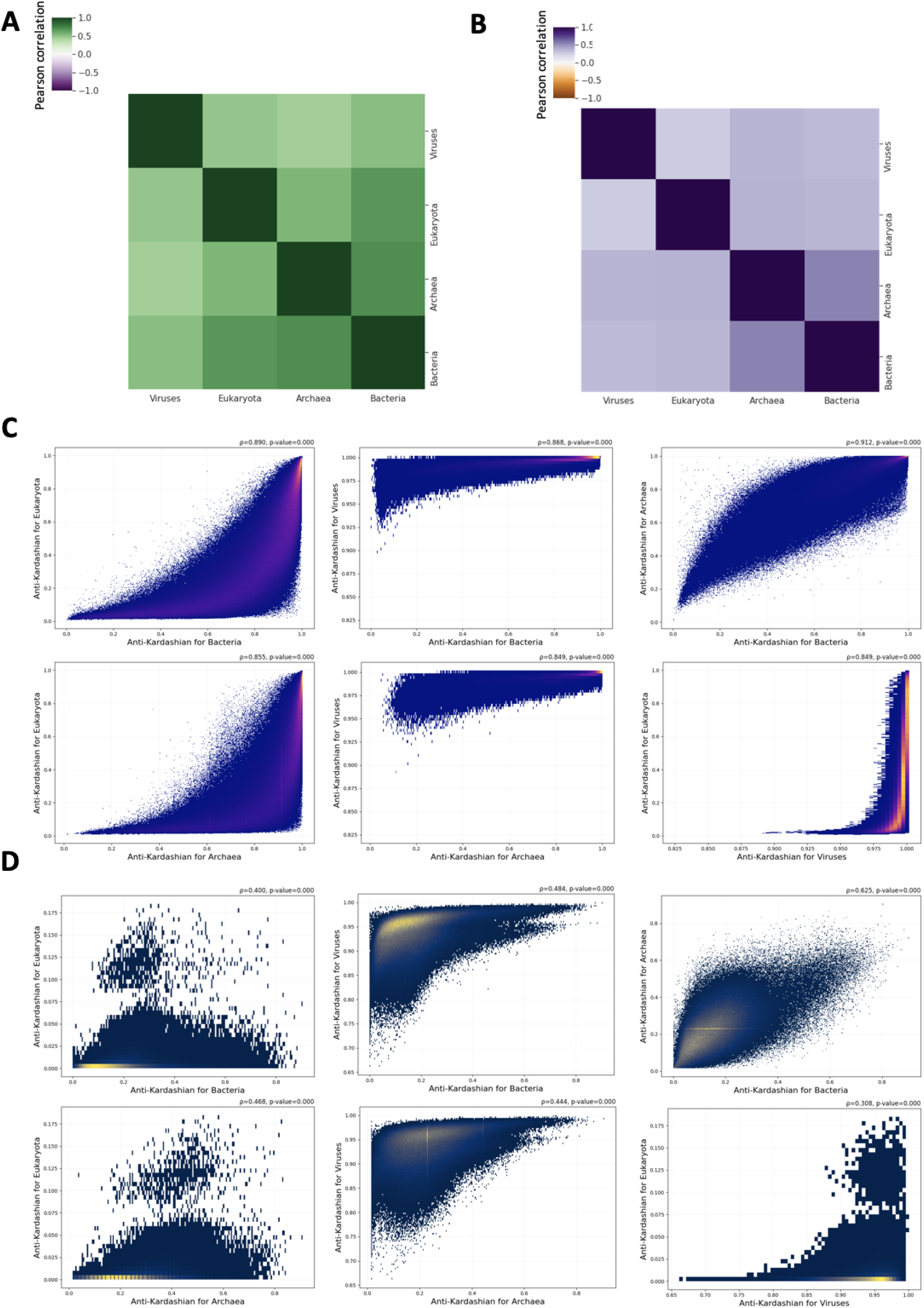
The Anti-Kardashian index between taxonomic sub-groups for peptide and nucleic kmers. **A**. Pearson correlation matrix of the Anti-Kardashian index between viruses, bacteria, archaea and eukaryotes for peptide kmers. **B**. Scatter-plot showing the correlation of the Anti-Kardashian index between pairs of taxonomic subgroups for peptide kmers. **C**. Pearson correlation matrix of the Anti-Kardashian index between viruses, bacteria, archaea and eukaryotes for nucleic kmers. Results shown for five amino acid kmers. **D**. Scatter-plot showing the correlation of the Anti-Kardashian index between pairs of taxonomic subgroups for nucleic kmers. Results shown for ten bp kmers. (**C-D**) Spearman correlation and associated p-values are shown.

We were also interested in investigating taxonomic differences in the amino acid and dinucleotide frequencies across the Anti-Kardashian spectrum, which could explain the differences in kmer rarity between the taxonomies. We therefore examined the amino acid and dinucleotide content of kmers within individual taxonomies as a function of the Anti-Kardashian index scores. For amino acids, we find surprisingly that in viruses, alanine and glycine are the primary amino acids found in commonly used kmers, whereas other amino acids are less likely to be shared between viral proteomes (**Supplementary Figure 8**). Bacteria, archaea and eukaryotes show similar profiles with tryptophan, methionine and cysteine being over-represented in kmers with the highest Anti-Kardashian index scores (**Supplementary Figure 8**). In contrast, peptide kmers with high Anti-Kardashian index scores in viruses do not show a preference for tryptophan, methionine or cysteine (**Supplementary Figure 8**). These compositional differences for rare kmers between viruses and other taxonomic groups account for the weaker correlation of the taxonomic Anti-Kardashian index in viruses with the other taxonomies (**Figure 4a**).

We examined nucleic kmers, for dinucleotide usage differences between taxonomies. We find that multiple dinucleotides show differences in their frequency profiles, including GC/CG, which are over-represented in eukaryotes relative to bacteria, for high Anti-Kardashian index (**Supplementary Figure 9**). In contrast, bacteria, archaea and viruses show higher levels of GG/CC at high Anti-Kardashian index scores relative to eukaryotes (**Supplementary Figure 9**). We also note that with the notable exception of viruses, the compositional differences between taxonomies for nucleic kmers with high Anti-Kardashian index are larger, than for peptide kmers.

### Sequence-based and physicochemical-based models of peptide kmer rarity in nature

Next, we investigated if we can predict the rarity of each peptide kmer in nature using as features its mono- and dipeptide sequence profile. Using ridge regression, we generated a model that predicts the rarity of peptide kmers, as measured by the Anti-Kardashian index, for peptide kmer lengths of three to six amino acids. The coefficient of determination (R^2^) of the model ranged between 0.972 and 0.561 for peptide kmer lengths of three to six amino acids, respectively (**Supplementary Figure 10**), indicating that the amino acid and dipeptide profile is largely predictive of the rarity of a peptide sequence in nature. We also examined the most informative coefficients of the ridge regression model. We observe that the most informative features are amino acids relative to dipeptides including Cysteine, Tryptophan, Alanine and Leucine, of which Alanine and Leucine are associated with a lower Anti-Kardashian index whereas Cysteine, Tryptophan are associated with a larger Anti-Kardashian index (**Supplementary Figure 11a-b**). These results are consistent with our earlier findings regarding differences in the amino acid composition of rare peptide kmers.

Nex, in order to capture non-linear relationships we developed a random forest regression model. We find that the model’s performance measured by R^2^ ranged between 0.968 and 0.816 for three and six amino acids kmer lengths (**Figure 5a-b**). When examining the feature importances in the model, we find amino acids to be more important in the predictive model than dipeptides across kmer lengths (**Figure 5e**; **Supplementary Figure 12**), Therefore, we conclude that the rarity of peptide kmers can be largely predicted using the amino acid and dipeptide content as features, with the amino acid composition of a peptide kmer providing the most significant predictive values.

**Figure 5:**
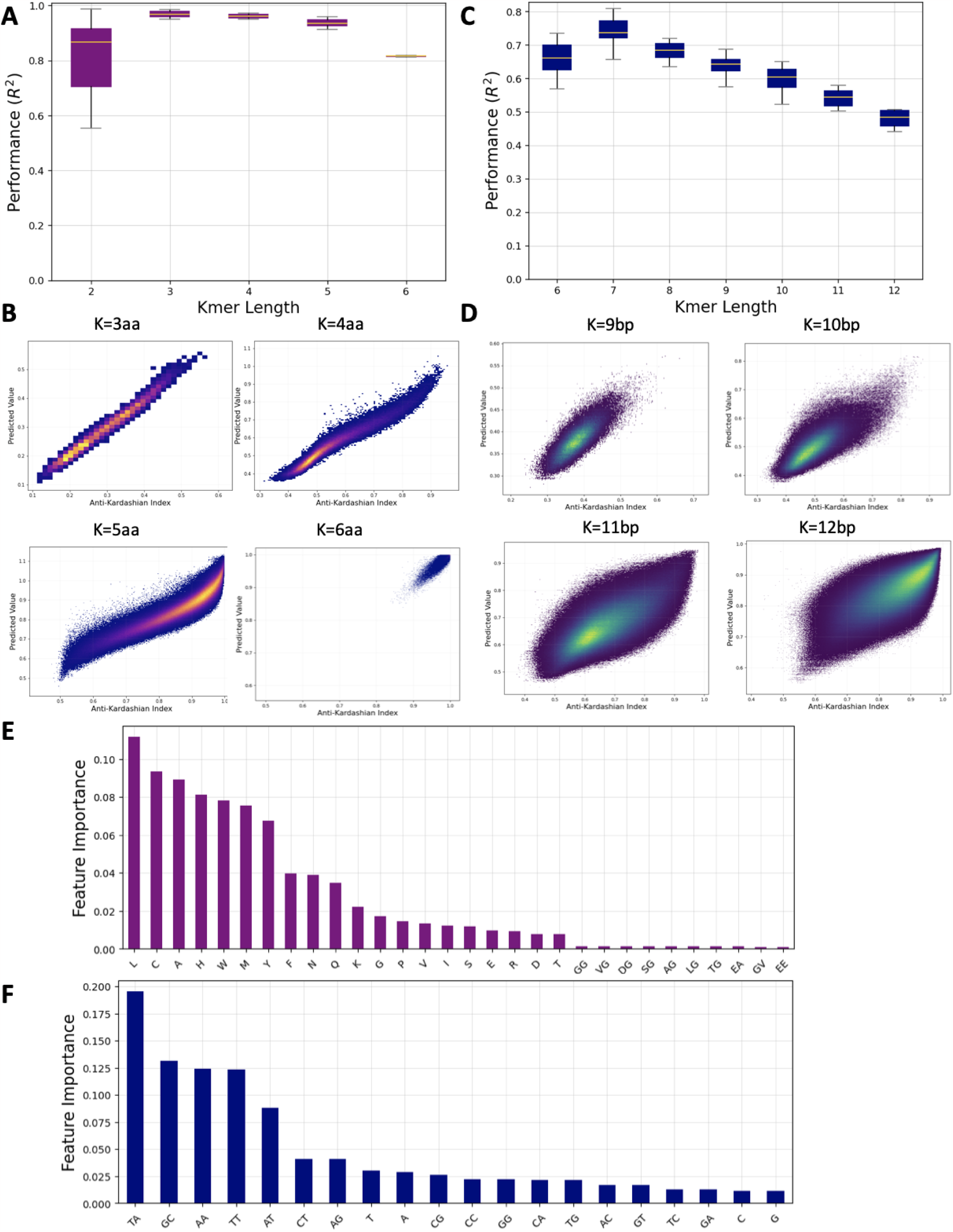
Predictive regression model determines the rarity of oligonucleotide sequences in nature. **A**. Regression model for the rarity of sixmer peptides. **B**. Regression model for the rarity of tenmer oligonucleotides. **C-D**. Coefficients of regression model for oligopeptides and oligonucleotides. **E**. Feature importances for the random forest regression model predicting the rarity of six amino acid peptides. **F**. Feature importances for the random forest regression model predicting the rarity of twelve bp nucleic kmers.

We examined if the physicochemical properties of the constituent amino acids of each peptide kmer can predict the Anti-Kardashian index score of each peptide kmer. The physicochemical features that were calculated for each kmer included the molecular weight, the isoelectric point, the charge, the hydrophobicity, the hydrophobic moment, the Boman index, the aliphatic index and the instability index. We next generated predictive models, based on these physicochemical properties of each oligopeptide. We report that the performance of a ridge regression model (R^2^) ranged in three-mers and six-mers between 0.426 and 0.200 (**Supplementary Figure 13a-b**). We observe that in contrast to the sequence-based model, a ridge regression has limited predictive power, when using the set of physicochemical features across the examined kmer lengths. Therefore, we also developed a random forest regression model that can capture non-linear relationships. We find that the random forest regression model trained on the physicochemical features achieves comparable results to the sequence-based model, with R^2^ ranging between 0.950 and 0.788 for kmer lengths of three and six amino acids (**Supplementary Figure 13c-d**). We also note that physicochemical properties of peptides are linked to their amino acid profile and are correlated features. We conclude that sequence-based and physicochemical-based regression models are predictive of peptide kmer rarity in nature and capture most of the variance between organismal proteomes.

### Sequence-based models of nucleic kmer rarity in nature

We were interested to examine whether there are sequence-based features that can be useful in predicting the rarity of nucleic sequences. To that end, we developed a sequence-based model that can predict the rarity of oligonucleotide kmer sequences across the 45,975 reference genomes studied. As features, we used the mononucleotide and dinucleotide profiles for each oligonucleotide kmer sequence of length between six and twelve bps. We first implemented a ridge regression model, to capture linear relationships, which was trained and tested separately for each kmer length. The model achieved R^2^ performance between 0.331 and 0.561, in six and twelve bp kmer lengths, respectively (**Supplementary Figure 10c-d**). As a result, we conclude that a linear regression model based on mono- and dinucleotide features cannot accurately predict the rarity of nucleic kmer sequences across organismal genomes.

To capture non-linear relationships, we used a random forest regression model, which significantly improved the performance of our model, across the kmer lengths studied. In particular, the model achieved R^2^ between 0.67 and 0.48, in six and twelve bp kmer lengths, outperforming the ridge regression model by up to 55% (**Figure 5c, d**). These findings indicate that the proportion of species in which nucleic sequences between six and twelve bps are found can be inferred to a significant extent from the mono and dinucleotide composition of these sequences.

Importantly, we examined the relative importance of each mononucleotide and dinucleotide in the performance of the random forest regression model. We report that the dinucleotides TA, GC and TT/AA are the most informative features in predicting the rarity of twelver-mer sequences in our model (**Figure 5f**; **Supplementary Figure 11**; **Supplementary Figure 14**). In contrast, G and C mononucleotides and GA/TC dinucleotides are the least informative features in our models (**Figure 5f**; **Supplementary Figure 11**; **Supplementary Figure 14**). These results could reflect the significance of the rarity of the TA dinucleotide across most organismal genomes which has been previously reported (Burge, Campbell, and Karlin 1992) and the influence of repetitive sequences which can be captured in repetitions of TT/AA motifs.

## Discussion

In this work, we have described the Anti-Kardashian index, a measure of the rarity of kmers, including oligopeptides and oligonucleotides, across organisms in natural sequences. In total, we analyzed 45,785 reference genomes and 21,871 reference proteomes and developed predictive models that can infer the rarity of a given sequence across the examined species. For oligopeptides, the amino acid composition of a kmer is the primary determinant of its rarity in nature, whereas for oligonucleotides we observe that its dinucleotide composition can partially account for its rarity. For peptide sequences, we also develop predictive models based on the biochemical properties of each oligopeptide sequence, and the performance of these models is close to those of the sequence-based models.

For peptide kmers, differences in amino acid molecular weight, energy expenditure, acquisition or synthesis costs influence the oligopeptide frequency spectrum in each organism (Swire 2007; Seligmann 2003; Akashi and Gojobori 2002) and likely account for the high predictive power of our models. For nucleic kmers, the kmer frequency spectra show differing distributions between species, including bimodal and unimodal spectra (Chor et al. 2009), and could account for differences in the Anti-Kardashian index scores when comparing different taxonomies. Dinucleotide compositional heterogeneity has been previously described across multiple species. Findings include the rarity of the TA dinucleotide across most organismal genomes (Burge, Campbell, and Karlin 1992) and the rarity of CG dinucleotides in vertebrates (Karlin, Campbell, and Mrázek 1998), both of which are consistent with our findings.

The Anti-Kardashian index can provide insights in evolutionary relationships between species, as it enables examining the set of kmers that are rare and only shared in a minority of organisms. In future work, we would be interested in examining other, finer taxonomic subdivisions, including potentially kingdoms and phyla. Indeed, we find significant differences between the three domains of life and viruses, which likely reflect differences in operative processes and selection constraints that shape their genomes and proteomes. Further work is also required to study the potential functions of rare and taxonomically biased kmers and their potential functional effects within evolutionary lineages.

The Anti-Kardashian index could also have multiple practical applications including in the development of biomarkers for the detection of pathogens and in wildlife conservation, by selecting rare kmers that are highly informative. Genetic engineering applications such as those using CRISPR could also benefit by selecting targets found in only a subset of the species. It will also be of interest to use the Anti-Kardashian index in metagenomic data and genome scaffolds. Finally, as more reference genomes and proteomes become available, the Anti-Kardashian index can be updated to more thoroughly reflect the rarity of kmers in nature. In summary, our work provides an index that estimates the rarity of each kmer in DNA and amino acid sequences and predictive models indicate that we can largely estimate the rarity of kmer sequences.

## Methods

### Reference genomes and proteomes used

Collection of reference genomes was performed for the genbank and Refseq databases as well as 104 reference genomes from the UCSC genome browser website. The genbank database included 26,854 bacteria, 438 archaea, 41,986 viral, 164 fungal, 5 plant and 1 invertebrate complete genomes. The Refseq database included 23,120 bacteria, 388 archaea, 11,128 viral, 19 fungal, 3 plant and 1 invertebrate complete genomes. Reference proteomes were downloaded from UniProt (https://ftp.uniprot.org/pub/databases/uniprot/current_release/knowledgebase/reference_proteomes/Reference_Proteomes_2022_03.tar.gz, Release 2022_03, 19-Sep-2022) and included a total of 344 archaea, 8623 bacteria, 2119 eukaryota and 10 789 viral reference proteomes. The total number of reference proteomes examined totalled 21,875 species.

### Derivation of the kmer profile of each genome and proteome

Identification of kmers was performed as previously described in (Mouratidis et al. 2023) for each genome and each proteome. For genomes, kmer lengths between six and twelve bps were used. For proteomes, kmer lengths of two to six amino acids were used.

### Calculation of the anti-Kardashian index

The number of species each kmer was found in was calculated. Based on this, the proportion of species a kmer was not found in was calculated from which we derived the anti-Kardashian index (see **Equation 1**). Similarly, the anti-Kardashian index was also calculated within available species of individual taxonomies, namely viruses, bacteria, eukaryotes and archaea. The frequency of each amino acid and each dinucleotide was calculated as a function of the Anti-Kardashian index in proteomes and genomes respectively. Clustering of species based on their kmer Anti-Kardashian profile was performed using the “clustermap” function of the Seaborn Python package, with default clustering parameters (Waskom 2021).

### Calculation of biochemical properties of peptides

For each peptide the following biochemical properties were calculated using the Peptides package (Osorio, Rondón-Villarreal, and Torres 2015): the Boman index, the Aliphatic index, the Instability index, its Hydrophobicity, the Hydrophobic moment, the isoelectric Point and its molecular weight.

### Oligopeptide regression models

The oligo-peptide sequence-based predictive model used as feature space the amino acid and dipeptide content of each oligopeptide. The oligo-peptide biochemical-based predictive model used as feature space the biochemical peptides. Ridge regression (l2 regularization) with strength regularization of alpha=0.01 was used from the Scikit-learn package (Pedregosa et al. 2012) using either the sequence-based features or the biochemical-based features. For each of the regression models, the coefficient scores of each feature were estimated from which the most predictive features were identified. Random forest regression was implemented to capture nonlinear patterns for the biochemical-based model and for the sequence-based model using the Scikit-learn package (Pedregosa et al. 2012). For determining feature importance the instance attribute feature_importances_ was used. Ten-fold cross-validation was performed for all models.

### Oligonucleotide regression models

The oligo-nucleotide sequence-based predictive model used as feature space the mono and dinucleotide content of each kmer. For a model that captures linear relationships, we used a ridge regression (l2 regularization) with strength regularization alpha=0.01 was used from the Scikit-learn package (Pedregosa et al. 2012). To capture non-linear relationships, we implemented a random forest regression model from the Scikit-learn package (Pedregosa et al. 2012) and the instance attribute feature_importances_ was used to derive the importance of each feature in the models. Ten-fold cross-validation was performed for all models.

## Acknowledgements

This study was funded by the startup funds of I.G.S. from the Penn State College of Medicine.

## Contributions

N.C., I.M., and I.G.S. conceived the study. I.G.S. supervised the study. N.C., M.M., and I.G.S. wrote the code, performed the analyses and generated the figures N.C. and I.G.S. wrote the manuscript with help from I.M, M.A.K., and A.M.

