## Supplementary Material for "The determinants of the rarity of nucleic and peptide short sequences in nature"

A

B

C

D

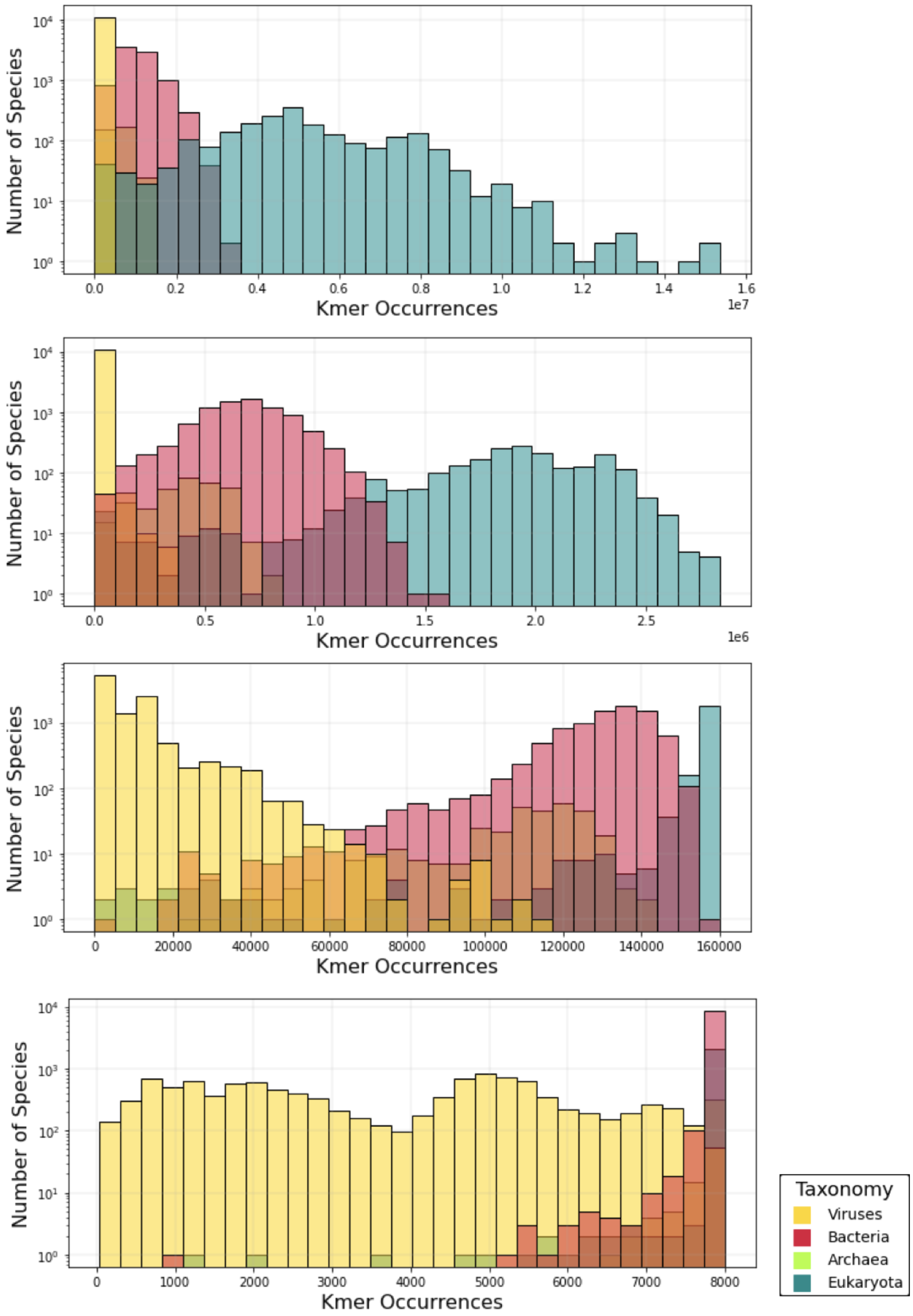


**Supplementary Figure 1: Number of species each peptide kmer was identified in.** Results shown for peptide kmer lengths of: **A.** six, **B.** five, **C.** four, **D.** three amino acids.

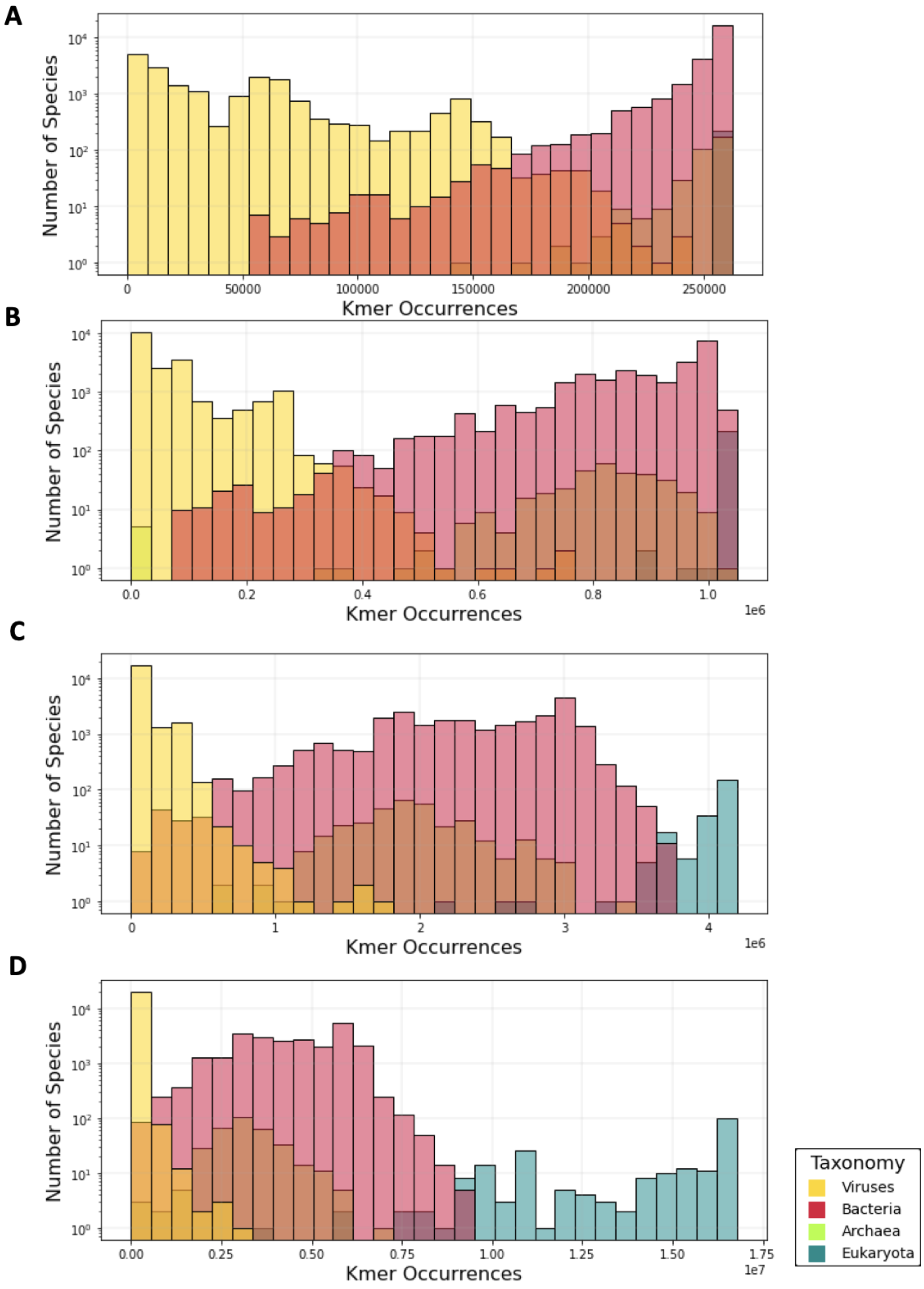


**Supplementary Figure 2: Number of species each nucleic kmer was identified in.** Results shown for nucleic kmer lengths of: **A.** nine, **B.** ten, **C.** eleven, **D.** twelve bps.

**A**

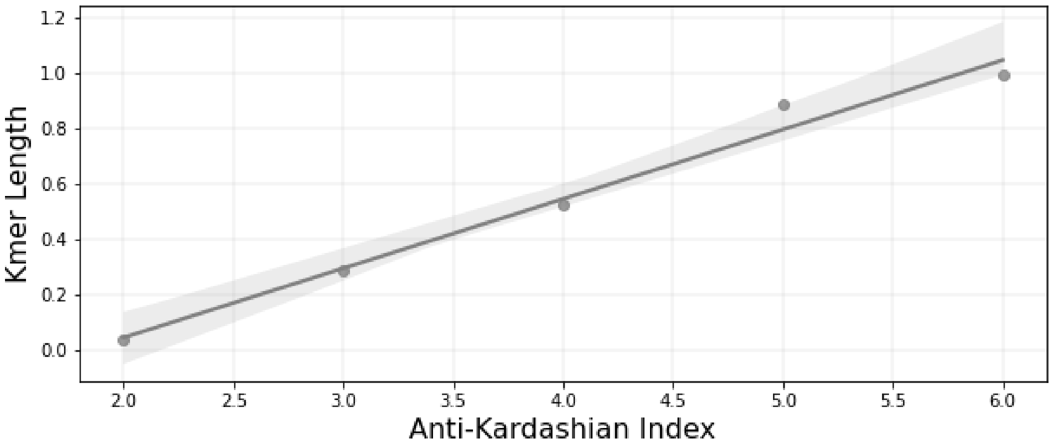


**B**

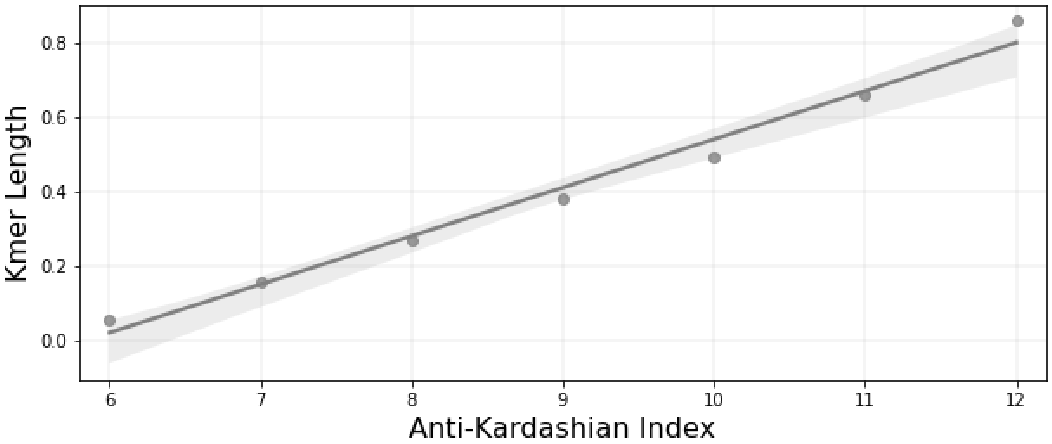


**Supplementary Figure 3: Association of Anti-Kardashian Index and kmer length for A. peptide kmers, B. nucleic kmers.** Error lines show 99 percentile confidence intervals.

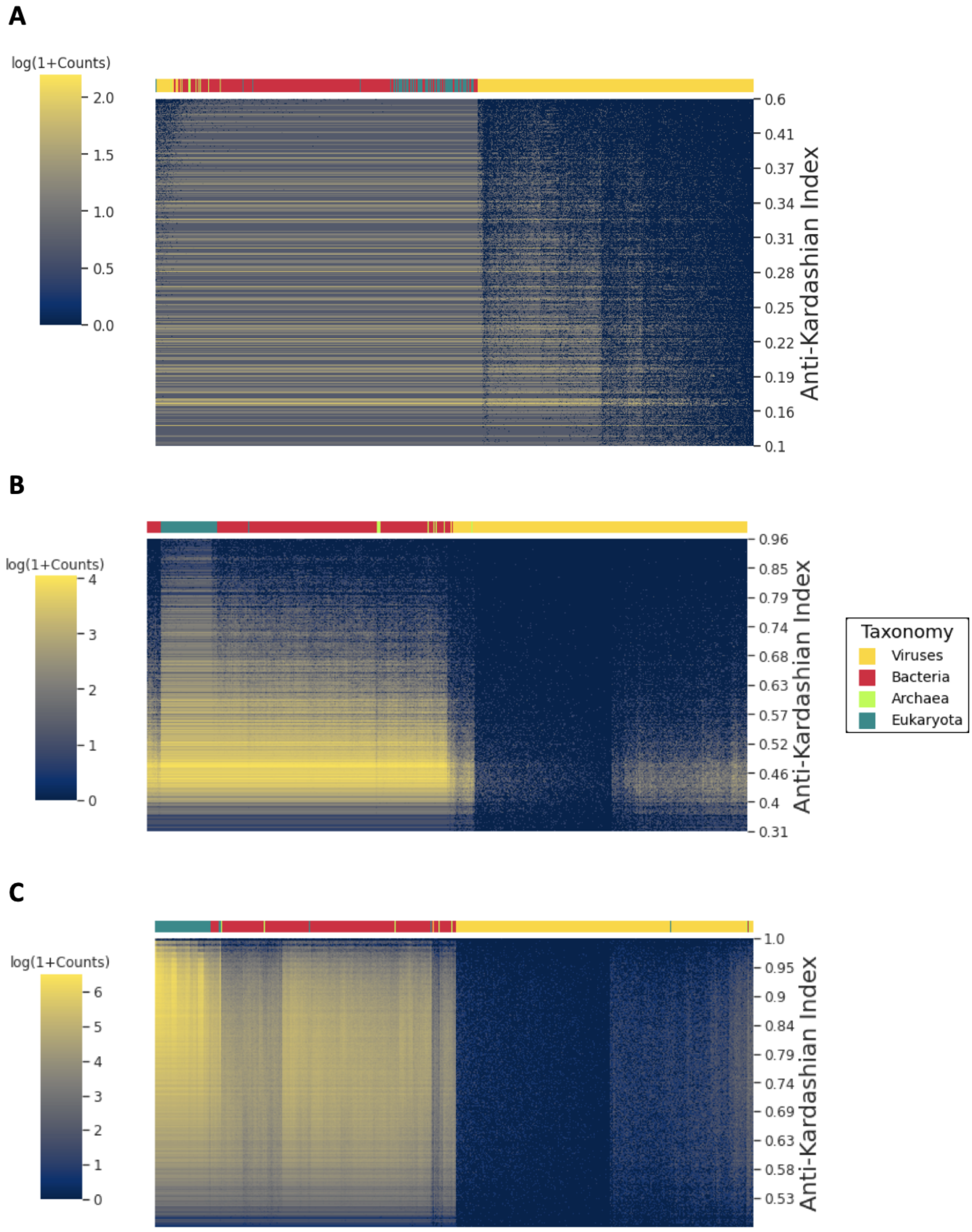


**Supplementary Figure 4: The number of kmers detected in each species as a function of the Anti-Kardashian index score.** Results shown for: A. three, B. four, C. five amino acids peptide kmer length. Color bar indicates the taxonomic group, namely viruses, bacteria, archaea and eukaryotes.

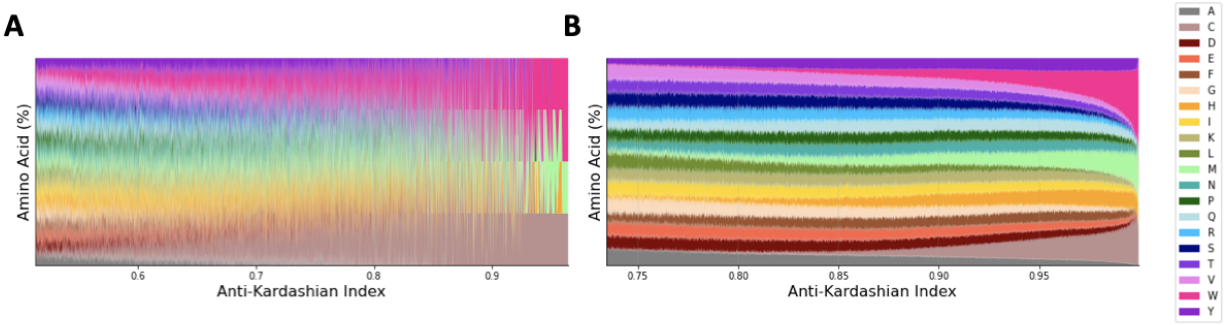


**Supplementary Figure 5: Association between the Anti-Kardashian Index and amino acid content of kmers in proteomes.** Results shown for: **A.** 4 amino acids, **B.** 5 amino acids.

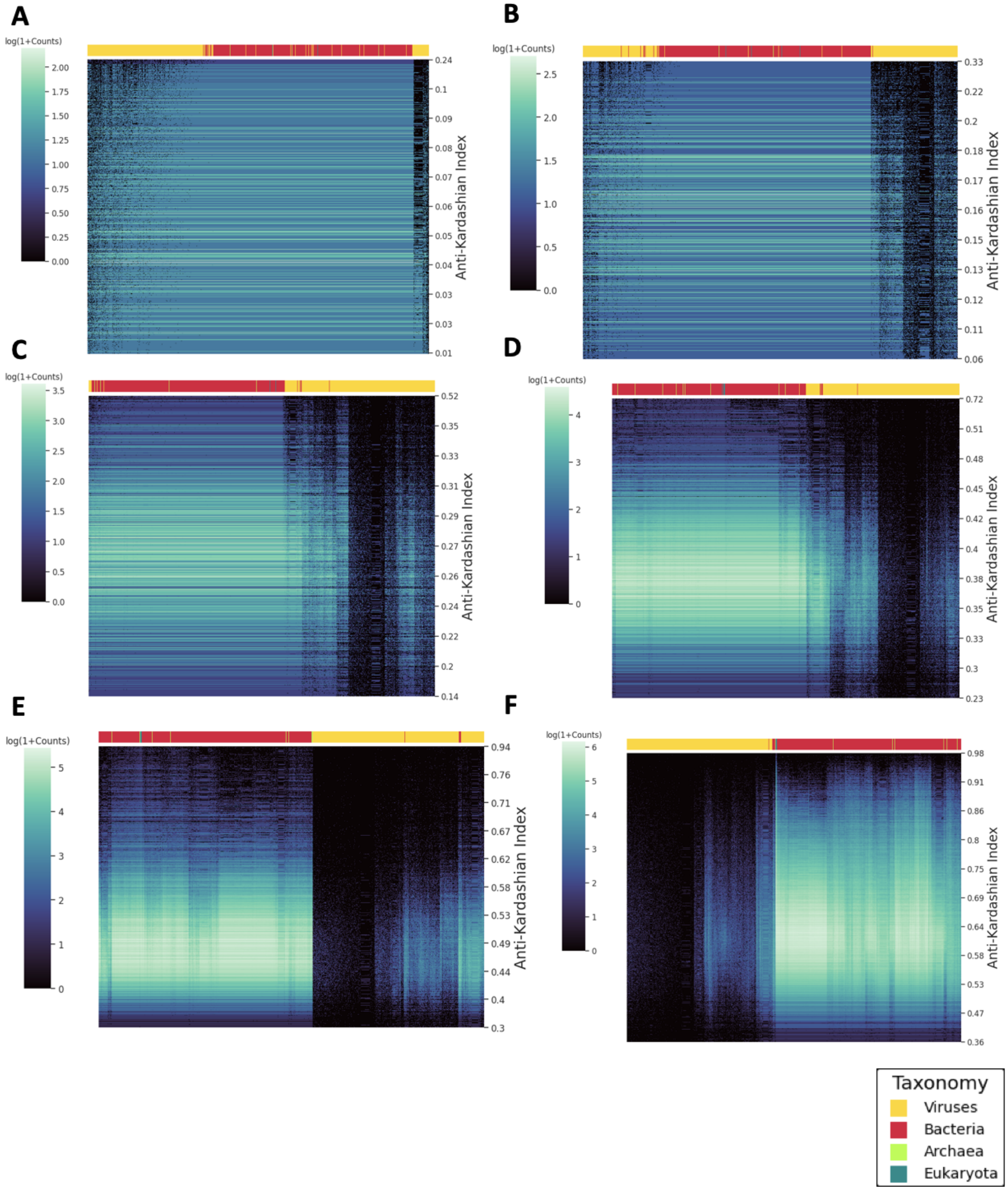


**Supplementary Figure 6: The number of nucleic kmers detected in each species as a function of the Anti-Kardashian index score.** Results shown for: **A.** six, **B.** seven, **C.** eight, **D.** nine, **E.** ten, **F.** eleven bps kmer length. Color bar indicates the taxonomic group, namely viruses, bacteria, archaea and eukaryotes.

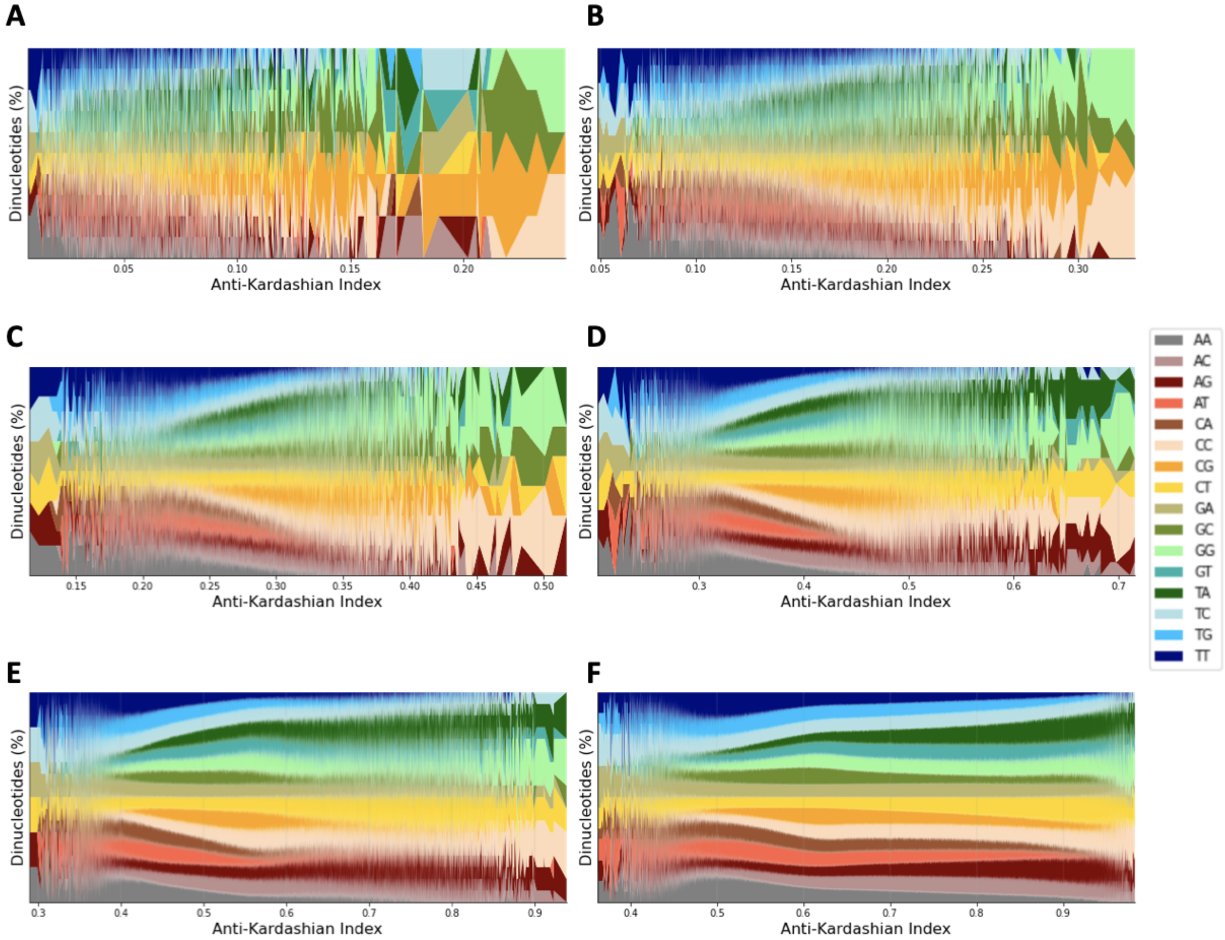


**Supplementary Figure 7: Association between the Anti-Kardashian Index and dinucleotide content of kmers in genomes.** Results shown for: **A.** 6bp, **B.** 7bp, **C.** 8bp, **D.** 9bp, **E.** 10bp and **F.** 11bp.

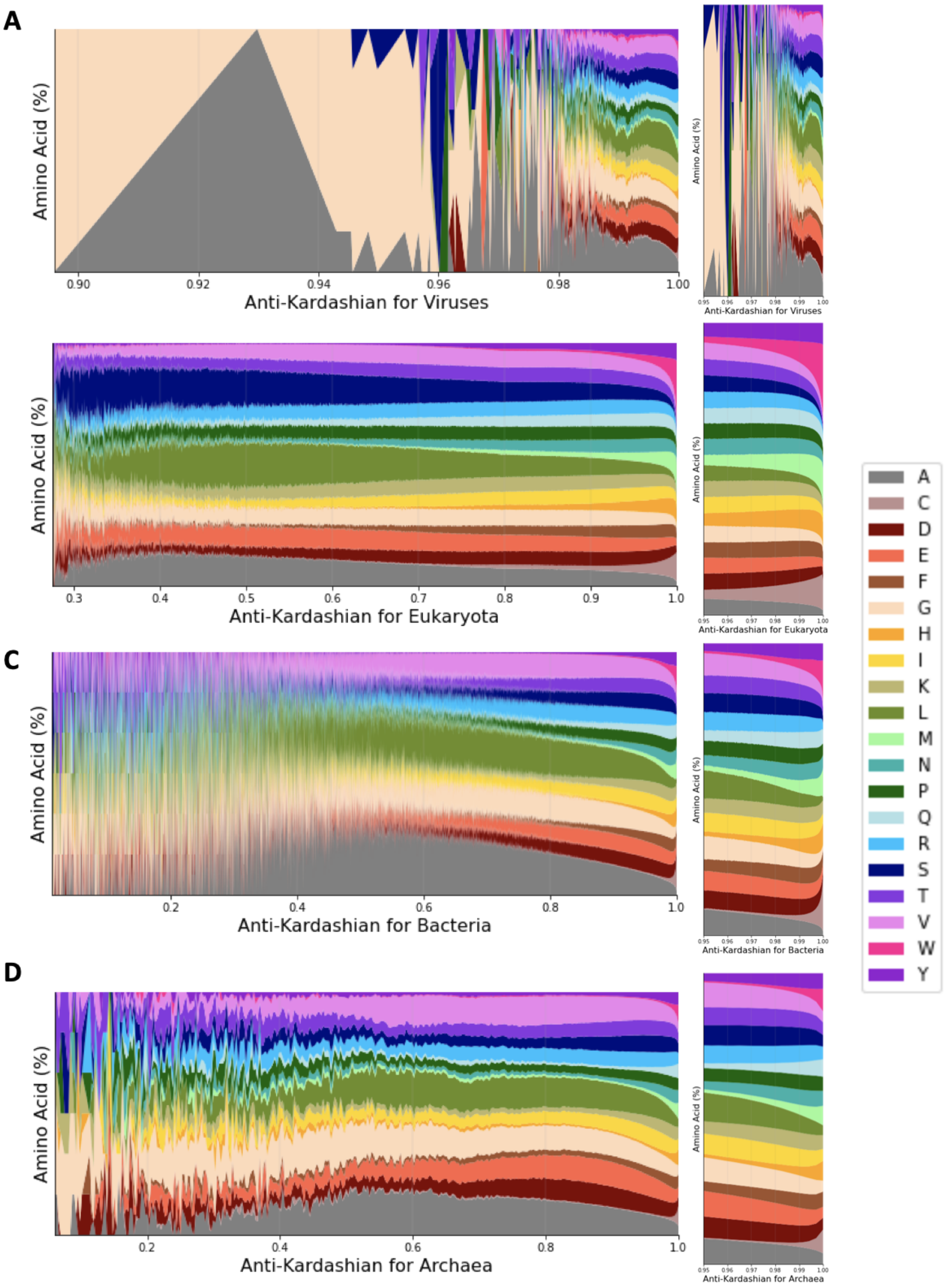


**Supplementary Figure 8: Association between the Anti-Kardashian Index and peptide content of kmers in proteomes.** Results shown for: **A.** Viruses, **B.** Eukaryotes, **C.** Bacteria and **D.** Archaea. Results shown for 6 amino acids kmer length.

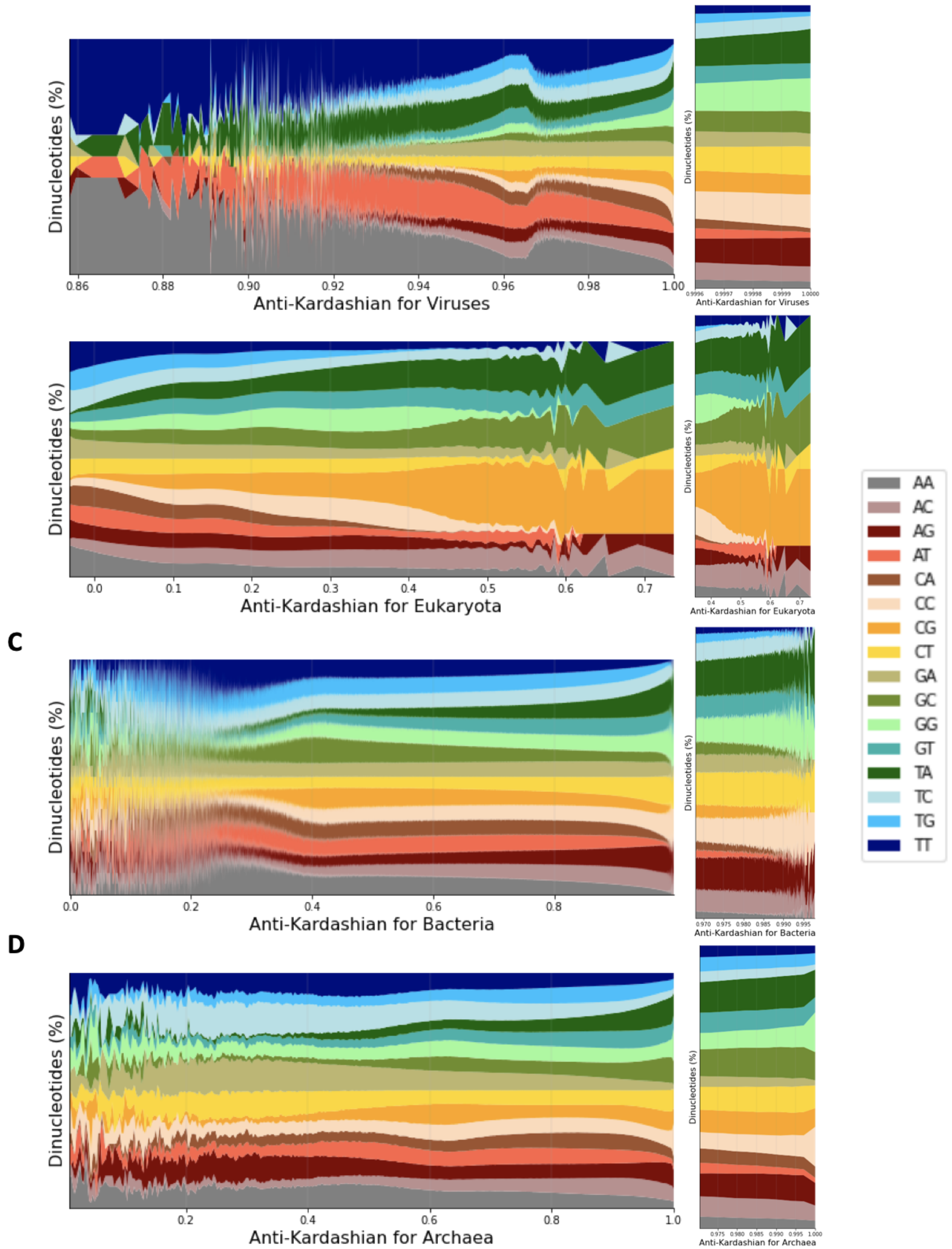


**Supplementary Figure 9: Association between the Anti-Kardashian Index and dinucleotide content of kmers in genomes.** Results shown for: **A.** Viruses, **B.** Eukaryotes, **C.** Bacteria and **D.** Archaea. Results shown for 12 bps kmer length.

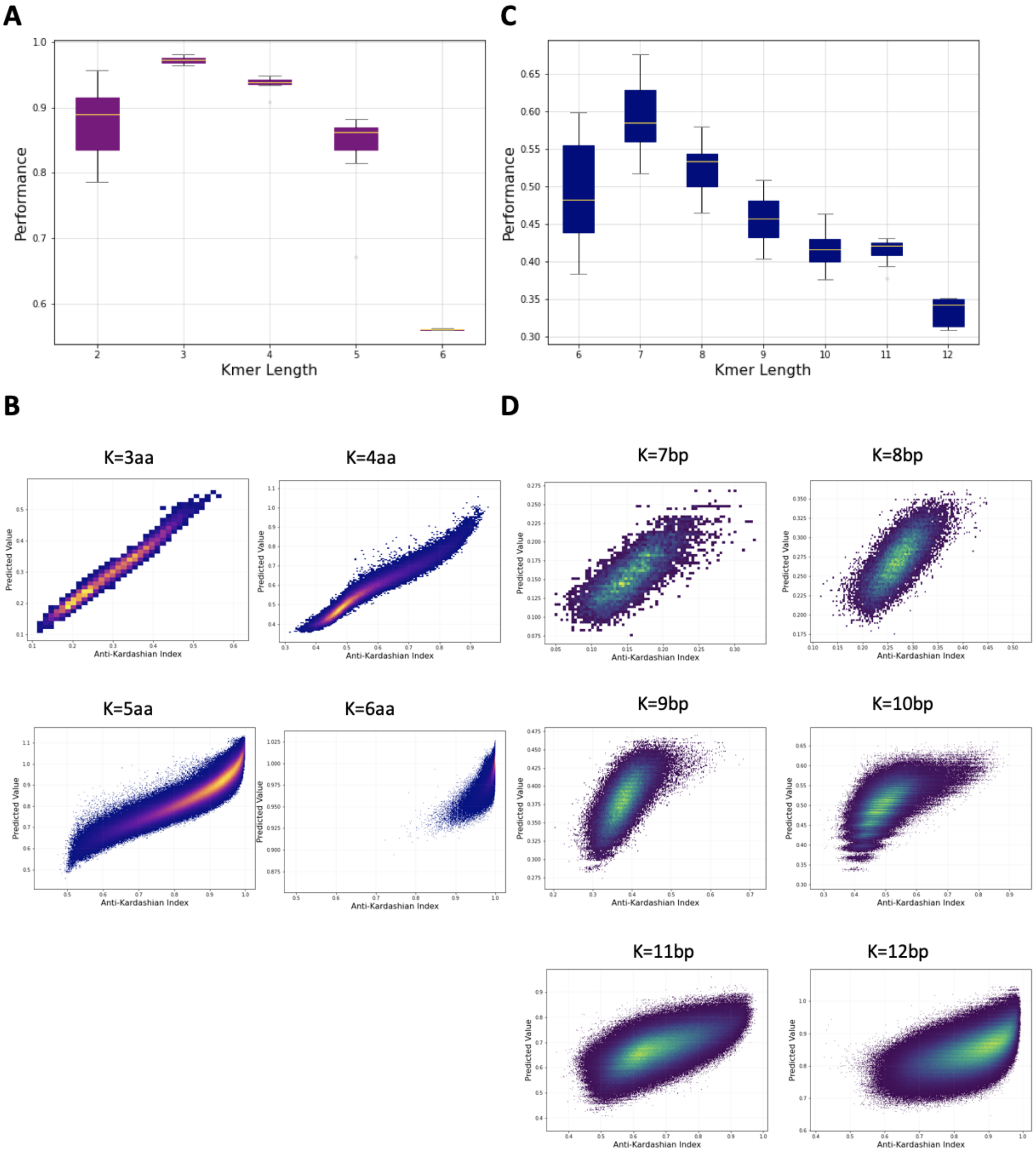


**Supplementary Figure 10: Predictive sequence-based models for the rarity of peptide and nucleic kmers across reference proteomes and genomes. A.** Performance of a ridge regression model for kmer lengths of six to twelve bp kmer lengths. **B.** Anti-Kardashian index versus predicted value for each kmer for kmer lengths of three to six amino acids (aa). **C.** Performance of a ridge regression model for kmer lengths of six to twelve bp kmer lengths. **D.** Anti-Kardashian index versus predicted value for each kmer for kmer lengths of seven to twelve bps.

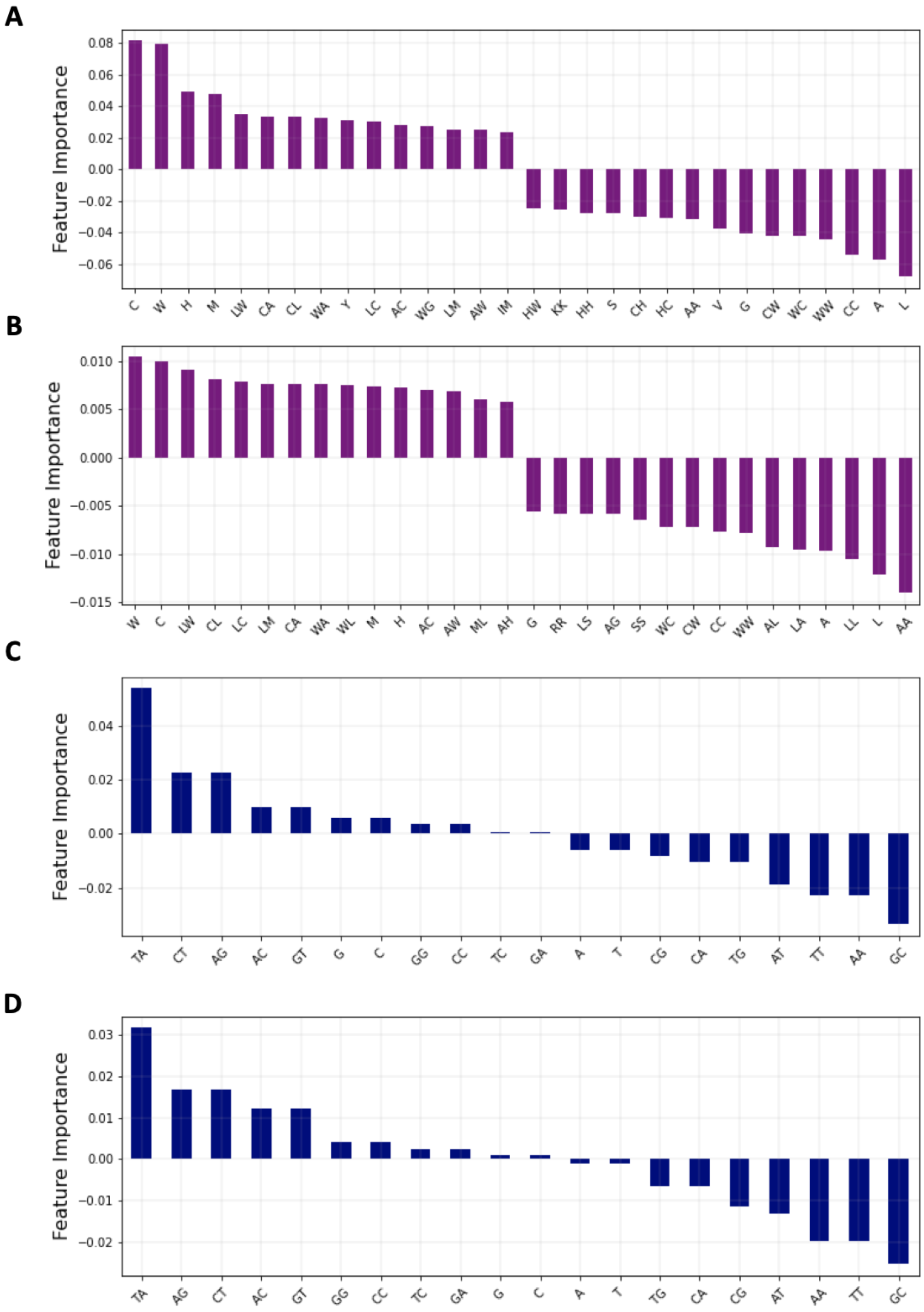


**Supplementary Figure 11: Feature importance for the coefficients of ridge regression models for peptide and kmer rarity.** Results shown for: **A.** 5 amino acid kmer length, **B.** 6

amino acid kmer length, **C.**11 bp kmer length, **D.**12 bp kmer length. The fifteen most positive and fifteen most negative coefficients are shown in A. and B.

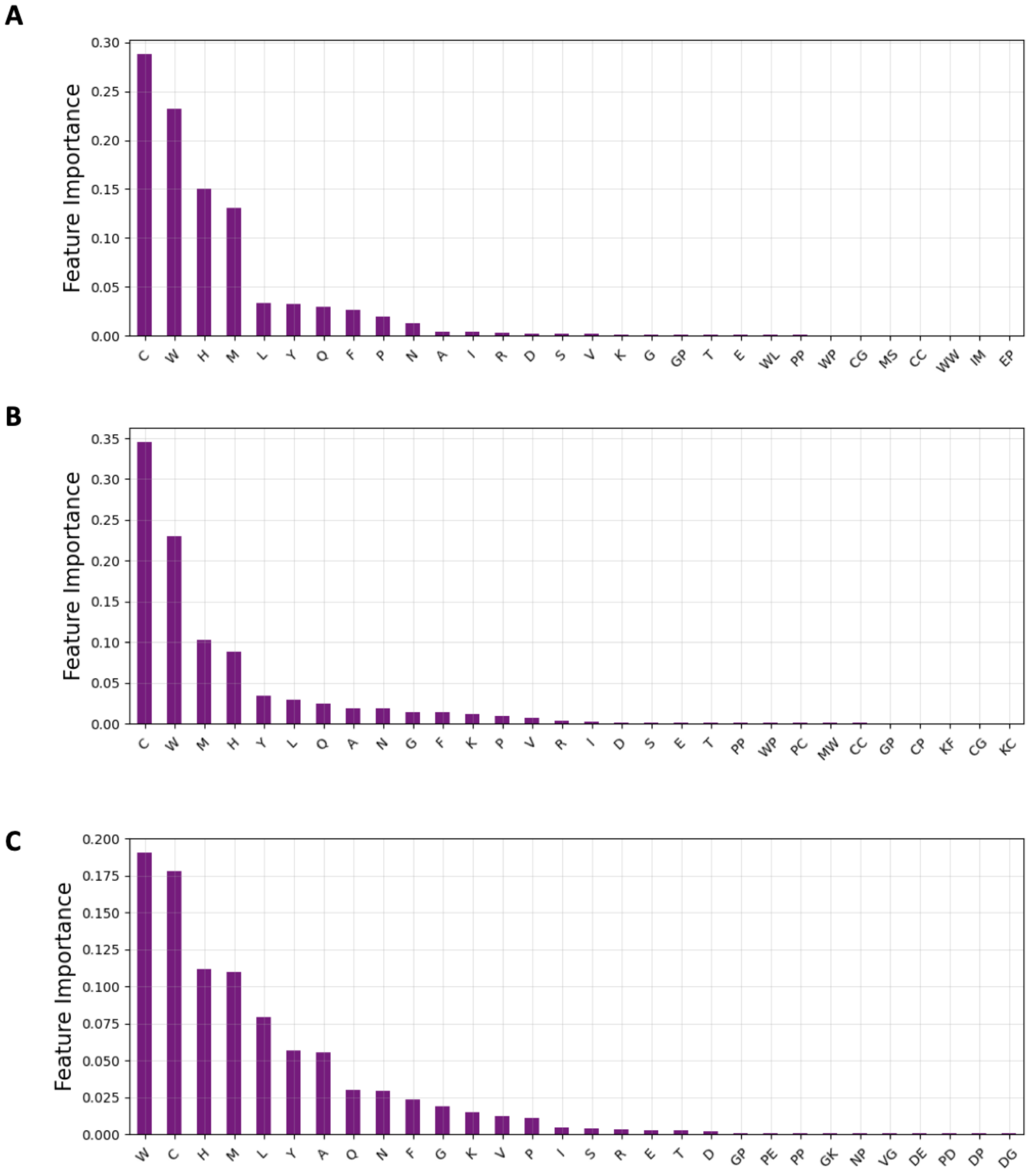


**Supplementary Figure 12: Feature importance for the coefficients of random forest regression models for peptide rarity.** Results shown for: **A.** 3 amino acid kmer length, **B.** 4 amino acid kmer length, **C.** 5 amino acid kmer length. The thirty most informative coefficients are shown in the panels.

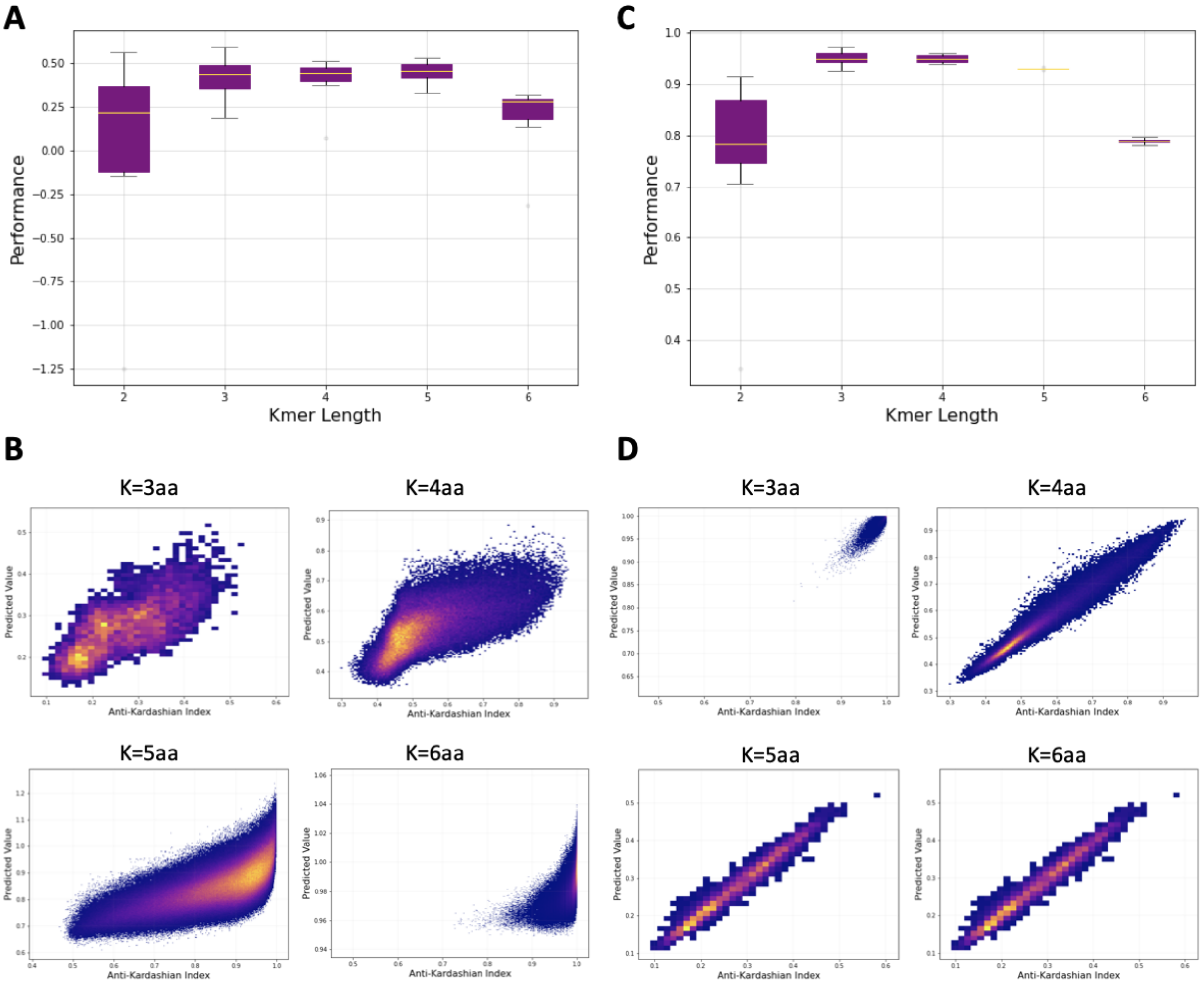


**Supplementary Figure 13: Predictive physicochemical-based models for the rarity of peptide kmers across organismal genomes. A.** Performance of a ridge regression model for peptide kmer lengths of three to six amino acids kmer lengths. **B.** Anti-Kardashian index versus predicted value for each peptide kmer for kmer lengths of three to six amino acids kmer lengths using the ridge regression model. **C.** Performance of a random forest regression model for peptide kmer lengths of three to six amino acids kmer lengths. **D.** Anti-Kardashian index versus predicted value for each peptide kmer for kmer lengths of three to six amino acids kmer lengths using the random forest regression model.

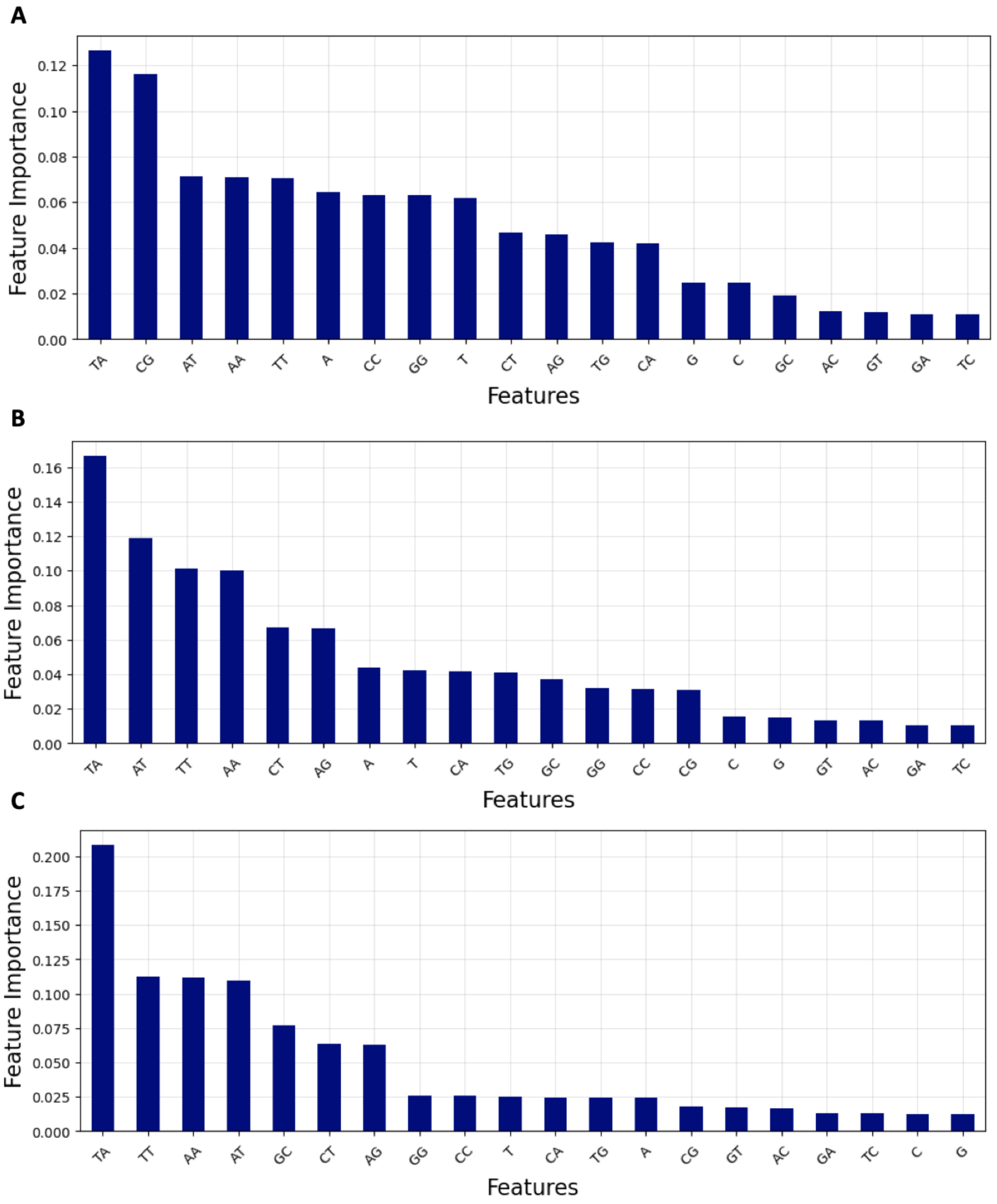


**Supplementary Figure 14: Feature importance for the coefficients of random forest regression models for nucleic rarity.** Results shown for: **A.** 9bp kmer length, **B.** 10bp kmer length, **C.** 11bp kmer length. The thirty most informative coefficients are shown in the panels.
